# Human 28S rRNA analysed by state-of-the-art oligonucleotide mass spectrometry: benchmarking current capabilities and a call to action for MS-Seq

**DOI:** 10.64898/2026.07.17.739151

**Authors:** Jannick Schicktanz, Yuyang Qi, Junzhou Wu, Bibek Hamal, Antony Lechner, Sam Wein, Oskar Knittelfelder, Nur Yesiltac-Tosun, Felix Dalwigk, Katharina Hoja, Leona Rusling, Sofia Obersteiner, Ella Bellenberg, Kira Kerkhoff, Michael S. DeMott, Robert Ross, Philippe Wolff, Kathrin Breuker, Patrick A. Limbach, Peter Dedon, Stefanie Kaiser

**Affiliations:** Goethe University Frankfurt, Institute of Pharmaceutical Chemistry, Max-von-Laue-Str. 9, 60438 Frankfurt, Germany; Antimicrobial Resistance Interdisciplinary Research Group, Singapore-MIT Alliance for Research and Technology Centre, 138602 Singapore; Department of Biological Engineering, Massachusetts Institute of Technology, Cambridge, MA 02139 USA; Rieveschl Laboratories for Mass Spectrometry, Department of Chemistry, University of Cincinnati, Cincinnati Ohio 45221 United States; Centre for Integrative Biology (CBI), Department of Integrated Structural Biology, IGBMC, 1 rue Laurent Fries, Illkirch, France; Centre National de la Recherche Scientifique (CNRS) UMR 7104, Illkirch, France; Institut National de la Santé et de la Recherche Médicale (Inserm) U964, Illkirch, France; Université de Strasbourg, Strasbourg, France; University of Tübingen, Institute for Bioinformatics and Medical Informatics, Maria-von-Linden-Str. 6, 72076 Tübingen Germany. OpenMS Inc. 502 W 7th ST, STE 100 Erie PA 16502 USA; Bruker Daltonics GmbH & Co. KG, Fahrenheitstr. 4, 28359 Bremen, Germany; Thermo Fisher Scientific, 10 Maguire Ave., Lexington, MA 01450, USA; Architecture et Réactivité de l’ARN, Institut de Biologie Moléculaire et Cellulaire, CNRS UPR9002, Université de Strasbourg, 2 allée Konrad Roentgen, Strasbourg, France; Institute of Organic Chemistry and Center for Molecular Biosciences Innsbruck (CMBI), University of Innsbruck, Innrain 80-82, 6020 Innsbruck, Austria

## Abstract

Oligonucleotide mass spectrometry (MS-Seq) is emerging as a powerful approach for sequence-resolved RNA modification analysis, yet the field lacks standards for experimental workflows, data analysis and reporting.

To assess current capabilities, the Human RNome Project Consortium conducted a cross-platform benchmarking study using a common RNA sample. A partial RNase T1 digest of human 28S rRNA was distributed to participating laboratories and analysed using existing LC-MS/MS workflows spanning different chromatographic strategies and mass spectrometers. To enable direct comparison, datasets were analysed using a harmonized NucleicAcidSearchEngine (NASE) workflow. Despite substantial methodological differences, laboratories recovered highly overlapping oligonucleotide sets and generated similar sequence coverage maps with a global coverage of 54.16%, demonstrating reproducible sequence information across platforms under standardized sample and analysis conditions. The benchmark further revealed incomplete sequence coverage, platform-specific differences in data architecture and increased assignment ambiguity during dynamic modification searches. Together with the community consensus developed during the HRPC workshop, these findings define priorities for the field, including improved sensitivity, standardized data analysis and reporting, community repositories, and robust bioinformatic workflows for confident de novo RNA modification discovery. This study provides an experimental benchmark and roadmap toward routine MS-based mapping of the human RNome.

**Graphical Abstract:** 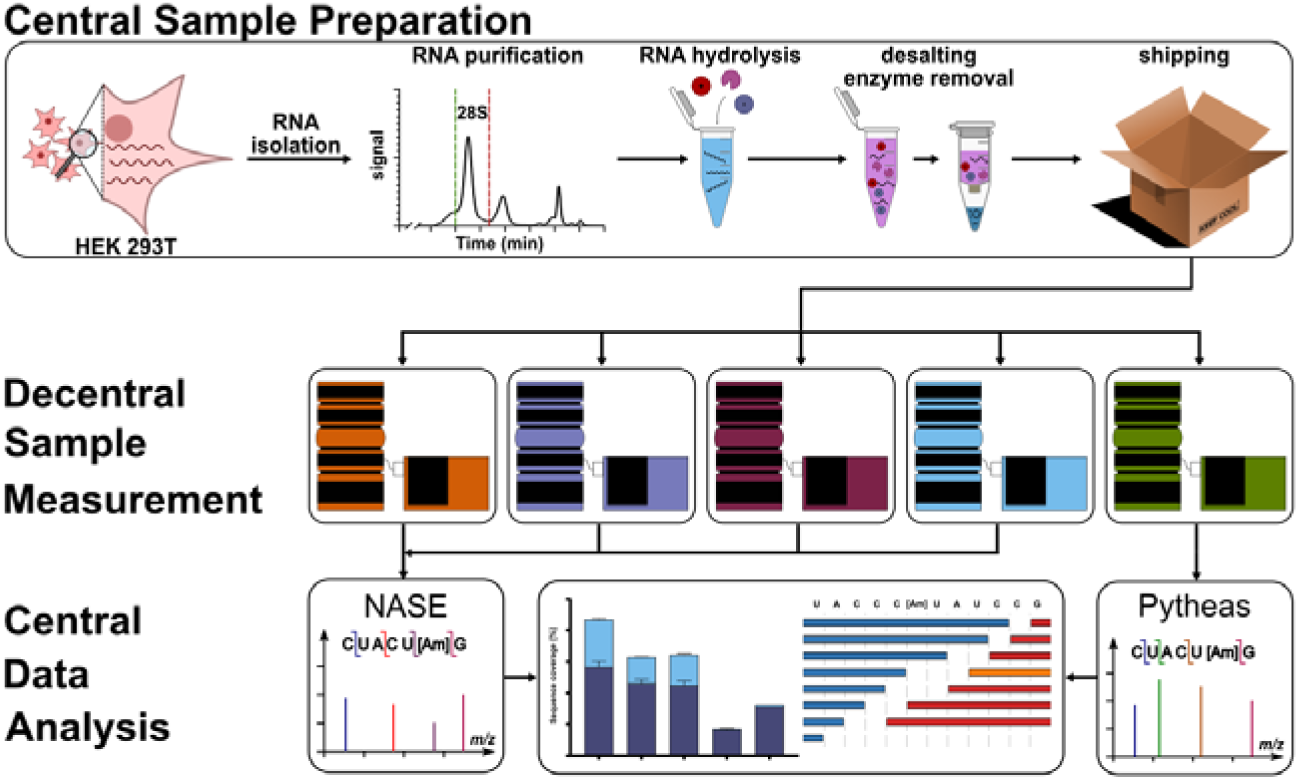

## Introduction

RNA modifications constitute an additional layer of gene regulation that expands the functional diversity of the transcriptome. More than 150 chemically distinct RNA modifications have been described to date across all domains of life and virtually every RNA class investigated (1). These modifications influence RNA folding, stability, localization, translation and interactions with proteins, and are increasingly implicated in human disease, therapeutic RNA design and cellular adaptation (2,3). Consequently, comprehensive characterization of RNA sequences together with their modification patterns has become a central objective in RNA biology.

Current analytical approaches largely rely on two complementary technologies: sequencing and mass spectrometry. Sequencing-based methods provide excellent sensitivity and transcriptome-wide coverage but generally infer RNA modifications indirectly through altered reverse transcription, nanopore current signatures or computational prediction models (4). By contrast, mass spectrometry directly measures the molecular mass of nucleotides and oligonucleotides and therefore provides orthogonal chemical information that is largely independent of prior modification knowledge (5). While nucleoside mass spectrometry has become a mature technology for absolute and relative quantification of RNA modifications (6), enzymatic hydrolysis to nucleosides inevitably destroys all sequence information and therefore prevents localization of modifications within individual RNA molecules.

To overcome this limitation, bottom-up oligonucleotide mass spectrometry has emerged as the method of choice for sequence-resolved analysis of modified RNAs (7–15). Similar to shotgun proteomics, RNA is digested into oligonucleotides (with e.g. RNase T1), separated by liquid chromatography, fragmented by tandem mass spectrometry and computationally reconstructed into sequence information. During the past decade, advances in chromatography, high-resolution mass spectrometry, and dedicated software such as NucleicAcidSearchEngine (NASE)(16) have transformed oligonucleotide MS from proof-of-principle studies into a broadly applicable technology capable of mapping RNA modifications in complex biological RNAs (14–17). Here we use the term *MS-Seq* throughout this manuscript to describe the comprehensive mass spectrometric analysis of RNA sequence and chemical modifications.

Despite these advances, MS-Seq remains far less standardized than proteomics (18–20). Laboratories employ diverse chromatographic approaches, mass spectrometers, fragmentation settings, RNases, database search strategies and performance metrics, making objective comparison between workflows difficult. Consequently, there is currently no consensus on how sequence coverage should be reported, how false discovery rates should be controlled, which metadata should accompany published datasets, or which analytical workflows are best suited for specific biological questions.

Community-wide benchmarking studies have played a pivotal role in establishing analytical standards in genomics (21), proteomics (19) and metabolomics (22) by objectively comparing technologies, defining quality metrics and identifying methodological bottlenecks. Such efforts not only accelerated technological development but also laid the foundation for community reference datasets and harmonized reporting standards that continue to support reproducible research. Except for the recent study focused on RNA modifications without any sequence context (22), there has been no comparable benchmarking study specifically focused on RNA sequencing by LC-MS.

To address this gap, the Human RNome Project Consortium (HRPC) initiated an international benchmarking effort bringing together laboratories with complementary expertise in RNA mass spectrometry. During the 2025 HRPC workshop, participating laboratories received identical aliquots of a ready-to-inject partial RNase T1 digest of human 28S rRNA, which they analysed using their established LC-MS workflows. By distributing a common sample while allowing laboratories to employ their preferred chromatographic, mass spectrometric and computational strategies, the benchmark was specifically designed to assess the current capabilities of state-of-the-art oligonucleotide mass spectrometry under standardized conditions. In parallel, the benchmarking results formed the basis for a community discussion on the future development of MS-Seq, resulting in the consensus recommendations presented in this report.

The objectives of this study were therefore twofold. First, we sought to evaluate the reproducibility, sequence coverage, and analytical performance of current oligonucleotide LC-MS workflows using a common reference digest analysed across multiple laboratories. Second, we aimed to identify the principal technological and computational bottlenecks currently limiting routine implementation of MS-Seq and to formulate a community roadmap for future method development, standardization and collaborative resource building.

## Materials & Methods

### Generation of the shared 28S rRNA hydrolysate

#### Cell culture

For this study, HEK 293T cells were incubated in Dulbecco’s modified Eagle’s medium (DMEM) D6546 high glucose supplemented with 10% non-dialyzed fetal bovine serum (FBS) and 0.584 g × L^−1^ L-glutamine. Cells were grown in T25 cell culture flasks (Art. No. TPP90026) (Faust Lab Science, Kamen, Germany) with 5 mL of the mentioned medium at 37 °C, 10% CO_2_ in an Eppendorf CellXpert C170i (Eppendorf, Hamburg, Germany). To maintain optimal growth conditions calls were split every 2-3 days. Handling was done under sterile conditions in a laminar air flow bench (HeraSafe 2025, Thermo Fisher Scientific, Waltham, MA, USA).

#### Cell lysis and total RNA extraction

HEK 293T cells were grown until 80% confluency, the medium removed and 1 mL of TRI reagent (Zymo Research Europe GmbH, Freiburg, Germany) added per T25 cell culture flask. The suspension was transferred to a new reaction tube and 200 µL of chloroform added. After vortexing the mixture was set aside at room temperature for 10 min to facilitate phase separation. The tube was centrifuged at 4°C and 12000 × g for 10 min. The upper phase was transferred to a fresh reaction tube and RNA precipitated using 600 µL 100% isopropanol. To ensure complete RNA precipitation the samples were stored at -20°C overnight.

After precipitation, samples were centrifuged for 45 minutes at 4°C and 12000 × g. The supernatant was removed and the pellet washed using 180 µL ice-cold 70% ethanol. Samples were centrifuged again for 10 minutes and washed an additional time, before centrifuging again for 10 minutes. Afterwards, the supernatant was removed and the reaction tube was placed with an open lid on the benchtop to dry for 5 minutes. Total RNA was reconstituted using 20 µL of ultrapure water and stored at -20°C.

#### Purification of 28S rRNA using SEC

28S rRNA was purified using an Agilent HPLC 1200 Series (Agilent Technoliogies, Santa Clara, Ca, USA) with an Agilent Bio SEC-5 1000Å (5 µm, 4.6 × 300 mm) column and column temperature of 40 °C. An isocratic elution with 0.1 M ammonium acetate (pH 7) at a flow rate of 1 mL × min^.1^ was used. Before running the separation, all samples were combined and the concentration measured using a Nanophotometer (N60, Implen, Munich, Germany). The volume for each injection was adjusted to contain 50 µg of RNA. After collecting the desired RNA fraction the volume of each sample was reduced to 50 µL using a Savant SpeedVac SPD120 (Thermo Fisher Scientific, Waltham, MA, USA). The RNA was precipitated using 0.1 × sample volume 5 M ammonium acetate and 2.5 × sample volume ice-cold 100% ethanol. To facilitate precipitation samples were incubated at -20°C overnight. The precipitated RNA was centrifuged for 45 min at 4°C and 12000 × g. Supernatant was discarded and the pellet washed one time with ice-cold 70% ethanol before centrifuging again for 10 min and discarding the supernatant. After drying for 5 min open on the benchtop the samples were resuspended in 10 µL ultra-pure water each.

#### Digestion of 28S rRNA using RNase T1

For the digest 33.3 U RNase T1 (Thermo Fisher Scientific, 1000 U/µL) per µg of 28S rRNA were used. The samples were digested in one batch in RNase T1 buffer with a final concentration of 25 mM TRIS (pH 7.5) and 100 mM NaCl. Digestion was carried out at 37°C for 30 min. Immediately after digestion, samples were purified using the Zymo Oligo Clean & Concentrator kit (C1004-50, Zymo Research Europe GmbH, Freiburg, Germany) according to the manufacturer’s protocol. The 330 µg RNA digest was separated into 10 µg aliquots each and the volume adjusted to 50 µL using ultra pure water prior to following the vendor’s protocol. Samples were eluted in 10 µL of ultrapure water, pooled and the concentration adjusted to 1000 ng/µL. 25 µg aliquots were prepared and stored at -80 °C until shipped. Shipping was performed in aqueous solution on dry ice, as recently described (6).

#### Polyacrylamide Gel Electrophoresis

Digested samples were analyzed using a 20% TBE-urea PAGE (prepared from ROTIPHORESE Sequencing gel concentrate, 3043.1, ROTIPHORESE Sequencing gel diluent, 3047.1 and ROTIPHORESE Sequencing gel buffer concentrate, 3050.1; Carl Roth, Karlsruhe, Germany). After polymerization the gel was transferred to a running chamber containing 1x TBE buffer, the gel was pre-run at 250V for 30 min. Samples and ladder (contained 76, 40, 30, 20, 9 and 5 nucleotides long oligonucleotides) were mixed with 2 × RNA loading dye (NEB, Ipswich, MA, USA). About 1 µg of RNA was loaded onto the gel. The gel was run at 275 V for 60–90 min and stained for 10 min with StainsAll solution (0.65× TBE, 0.01% StainsAll, 10% formamide, 25% isopropanol, 65% water). After staining the gel was destained for 2x 30 minutes destaining solution (1× TBE buffer with 25% isopropanol) and imaged using a Bio-Rad ChemiDoc™ MP Imaging System.

#### LC-MS/MS Workflows

Four laboratories contributed five LC-MS/MS workflows, hereafter referred to as systems 1-5. Below we provide a brief overview of the LC-MS instrumentation used in this benchmarking study. Detailed chromatographic conditions, gradient, mobile-phase composition, mass spectrometric parameters and data acquisition settings for each system are provided in the accompanying metadata sheet.

### System 1: Analytical flow HILIC LC-MS/MS

System 1 employed analytical hydrophilic interaction chromatography (HILIC) LC conditions (i.e., 300 µL min^-1^ flow rate) with an Orbitrap Exploris 240 mass spectrometer (Thermo Fisher Scientific) for analysis. Data were acquired in data-dependent acquisition (DDA) mode with collision-induced dissociation (CID) in the HCD collision cell. The analytical workflow has been described previously.

### System 2: Microflow RP LC-MS/MS

System 2 employed microflow reverse-phase (RP) LC conditions (i.e., 85 µL min^-1^ flow rate) with an Eclipse mass spectrometer (Thermo Fisher Scientific). Data were acquired in DDA mode with CID in the linear ion trap and orbitrap detection.

### System 3: Nanoflow HILIC LC-MS/MS

System 3 employed nanoflow HILIC LC conditions (i.e., 300 nL min^-1^ flow rate) with a Q Exactive Plus Orbitrap mass spectrometer (Thermo Fisher Scientific). Data were acquired in DDA mode with CID in the HCD collision cell. The analytical workflow has been described previously (15).

### System 4: Microflow IP-RP LC-MS/MS

System 4 employed microflow ion-pair reverse-phase (IP-RP) LC conditions (i.e., 5 µL min^-1^ flow rate) with a Synapt G2-S mass spectrometer (Waters). Data were acquired in DDA mode with CID in the Triwave collision cell. The analytical workflow has been described previously (23).

### System 5: Nanoflow IP-RP LC-MS/MS

System 5 employed nanoflow IP-RP LC conditions (i.e., 300 nL min^-1^ flow rate) with a Synapt G2-S mass spectrometer (Waters). Data were acquired in DDA mode with CID in the Triwave collision cell. The analytical workflow has been described previously (14).

### Conversion of vendor raw files into mzML files

The vendor specific raw files were converted to mzML files using MSConvert (https://github.com/ProteoWizard/, Version: 3.0.25229-e2d99c4). For all files, binary encoding precision was set to 64-bit, write index and TPP compatibility were enabled. Furthermore, a peak picking filter was applied for MS levels 1-2. For data from Waters instruments, the Waters DDA processing filter was added.

### NucleicAcidSearchEngine

The following parameters apply to both the static and dynamic search. Measurements from systems 1 – 4 were analyzed using NASE version 3.5.0 implemented in the OpenMS framework. For the target search all currently available human 28S rRNA reference sequences have been used (M11167.1, 7BHP_L5, RIBOMAP_HS28S, KROGH_HS28S, MOTORIN_HS28S, MARCHAND_HS28S). Here, the unmodified and modified references were used. For modified references, any phosphates and pseudouridines that were present in the sequence were removed. For decoy generation the decoy database was set to shuffle and RNA mode. For the NASE search, an FDR cut-off of 0.15 was used and all decoys removed. Precursor and fragment ion mass tolerance were set to 20 and 15 ppm, respectively. Charge states -1 to -20 were included as well as “unknown” charge states (referring to unassigned charge states). Adducts for precursor ions taken into consideration were sodium (Na^+^), potassium (K^+^), ammonia (NH_4_^+^) and acetate (C_2_H_4_O_2_^-^). The following isotopes were required to gain sequence information -1, 1, 2, 3 and 4. Oligo minimum size was set to 3 nucleotides and maximum size to 40. Unless indicated otherwise, one missed cleavage and RNase T1 was used. For all searches, all possible fragment ions were used (a-B, a, b, c, d, w, x, y, z). For the dynamic search only the addition of variable modifications was necessary. Here all modifications (apart from pseudouridine) that were present in the modified sequences were used, those include m^6^A, Am, m^1^A, m^5^C, nm^5^ges^2^U, m^3^U, Cm, Gm and Um. The number of variable modifications varied between 1 and 3 (exact parameters and configurations can be seen in Figures S6 and S7)

### Pytheas data analysis

For automatic data interpretation, LC-MS/MS raw files were converted to .mgf using PLGS (ProteinLynx Global Server, Waters Corp.) and ProteoWizard MSConvert. In silico digestion files were generated using Pytheas (23) with the corresponding RNA sequence and known modifications (pseudouridines were not searched). The converted .mgf files were processed using Pytheas to assign oligonucleotides. Low-score assignments (Sp < 0.1) were filtered out. Parameters used for data processing are provided in the accompanying Supplementary Information.

### Further data analysis of NASE and Pytheas data

Python scripts were created that specifically extract fragment information from generated result files.

To generate genome browser-style overlays and calculate global sequence coverage, the custom Python script 20260712_Genomebrowser_global_coverage.py was used. NASE .mzTab files and Pytheas final_report files were processed alongside their respective FASTA reference sequences. Global evaluation parameters were set to strictly filter for RNA cleavage after G (specific cleavage only), relying exclusively on NASE mzTab coordinates. For Pytheas data, tolerant mapping was disabled. Sample names were anonymized across datasets where necessary.

UpSet plots for the comparison of static search results were generated using the 20260712_Upset_plot_static_search.py script. The combined Excel report from the primary coverage analysis served as input, with plot intersections sorted by decreasing fragment count.

MS2 spectra and corresponding ion ladders were extracted and visualized using the 20260712_MS2_spectra_Ion_ladder.py script, integrating NASE and Pytheas results with their mzML files. For MS2 spectra visualization, the relative intensity threshold was set to 0%, charge annotations were hidden, and ghost-peak recovery was enabled. For the generation of fragmentation ladders, a compact layout was selected with a minimum intensity threshold of 5%.

To systematically compare static and dynamic NASE searches, the 20260712_Comparison_variable_static_search.py script (24) was applied. Both search results were evaluated against the modified reference sequence used for the static search. Mapping parameters were configured to strictly use G-specific cleavage and utilise mzTab coordinates.

The script 20260713_length_vs_RT.yp (24) was used to create chromatograms of the fragment length vs the retention time. Here the respective mzTab and final_report files were loaded and the parameters set to negative ion mode with yattering on the y-axis.

For all NASE analyses it is important to keep mzTab files in their original location as the script automatically searches for the corresponding mzML files by the relative path structure NASE uses.

All Python scripts, environment requirements, and detailed usage instructions are openly available in the associated GitHub repository (24).

### Use of AI language models

We used an AI assistant (GPT-5-5 Thinking) to support manuscript preparation in well-defined, author-directed tasks. Specifically, we prompted the model to support language editing for clarity, logical flow, structure, concision, terminology harmonization, and grammar. All AI-generated outputs were reviewed, edited, and verified by the authors; data analysis decisions, figures, and text content remain the responsibility of the authors. Googles Gemini 3.1 Pro and Anthropics Claude Sonnet 5 were used to support generation of python scripts for data analysis. All scripts and script outputs were manually curated and reviewed to ensure validity.

## Results

### Benchmark design and participating workflows

The workflow for RNA sequencing by mass spectrometry requires various steps starting from sample isolation and ending with data analysis. It is currently unclear how much variation each of these steps generate as no large-scale benchmarking study has yet been done for this technology platform. To better understand the effects of the analysis steps, all participating labs were provided identical samples of partially digested 28S rRNA prepared as described in the Materials & Methods. Supplemental Figure S1 shows the size distribution of the resulting hydrolysate to be in the range of 10-50 nucleotides. The hydrolysate was prepared for injection by desalting and 25 µg aliquots were analysed during the HRPC 2025 satellite workshop or shipped to participating labs afterwards (Figure 1A). It is noteworthy that the ready-to-inject hydrolysate gave reproducible results after storage at -20 °C for more than 3 months (Supplemental Figure S2).

**Figure 1.**
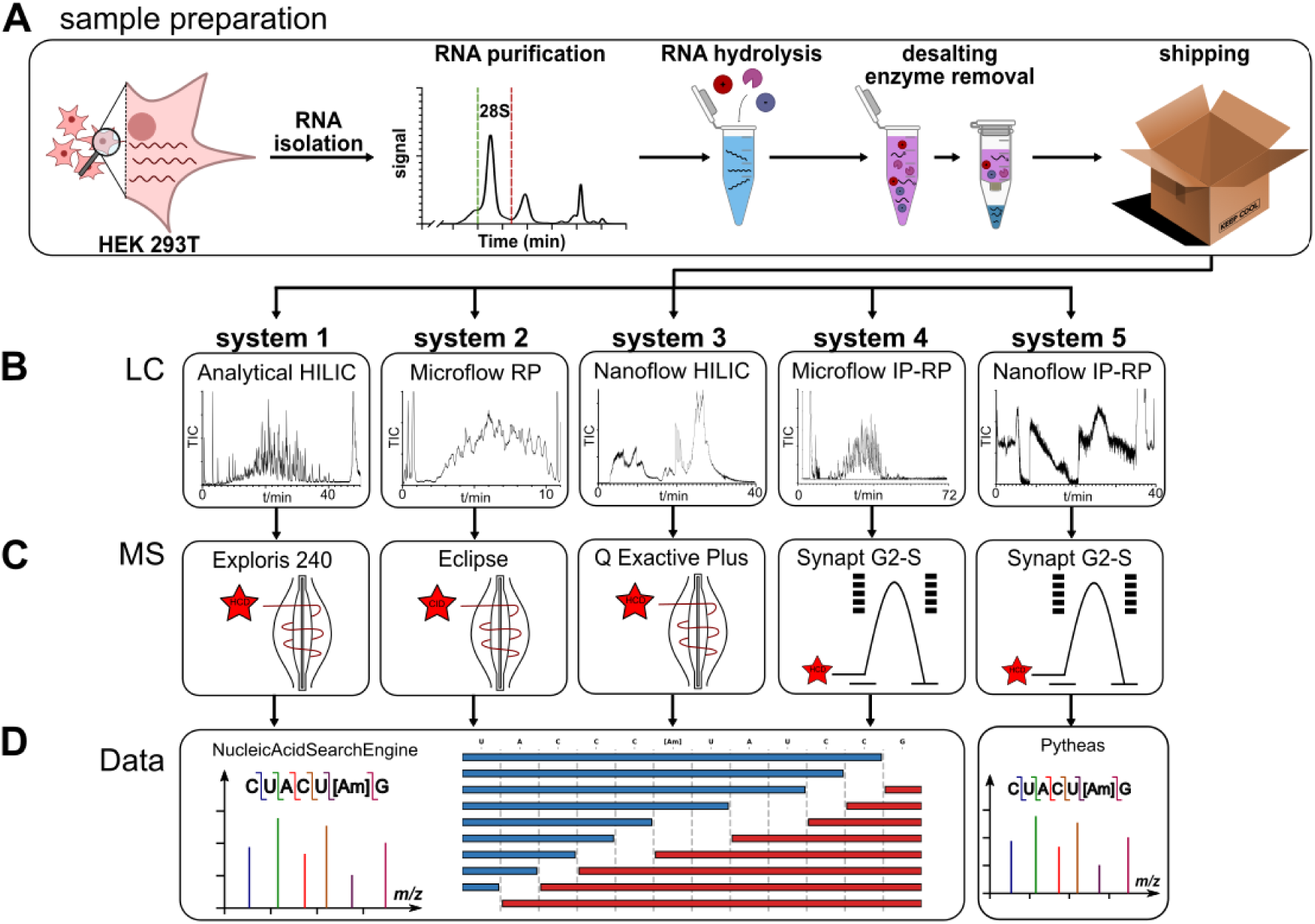
Experimental design of the HRPC oligonucleotide MS benchmarking study. **A** HEK 293T RNA was isolated and the 28S rRNA was purified by size exclusion chromatography. Following partial RNase T1 digestion, a common ready-to-inject oligonucleotide mixture was generated, quality controlled, aliquoted and distributed to the participants who analysed the digest using their established LC-MS/MS workflow. **B** chromatographic strategies and flow regimes used in this study. **C** Instrument platforms used in this study. **D** Raw data were subsequently analysed using either Pytheas (System 5 workflow) or a harmonized Nucleic Acid Search Engine (NASE) (all other workflows) to enable direct comparison of oligonucleotide identifications across laboratories.

Data was submitted from the five LC-MS/MS workflows used in these analyses (Table 1). Two datasets were acquired using HILIC, two using IP-RP, and one using RP (Supplemental Figure S3). For HILIC separation, the RNA partitions into the hydrophilic layer of the stationary phase due to its negatively charged phosphodiester backbone which causes a separation by length (15,25,26). RP separation is affected by interaction of the hydrophobic nucleobases with the stationary phase leading to retention based on base composition and hydrophobicity (13). The retention in RP can be advanced by addition of ion-pair reagents that mask the anionic phosphodiester backbone and cause separation of oligonucleotides mainly by length (14). Beyond the separation mode, three different flow rate regimes were used: two datasets were acquired using a nanoflow approach (300 and 350 nL/min), two datasets were acquired with a microflow approach (5 and 85 µL/min), and one with analytical flow (300 µL/min). An overview of reported sensitivities in this study is given in Supplemental Figure S4.

**Table 1:** Participating platforms in the 2025 HRPC workshop for MS-Seq benchmarking.

| Name | LC | MS |
| --- | --- | --- |
| System 1 | Analytical flow HILIC | Exploris 240 |
| System 2 | Microflow RP | Eclipse |
| System 3 | Nanoflow HILIC | Q Exactive Plus |
| System 4 | Microflow IP-RP | Synapt G2-S |
| System 5 | Nanoflow IP-RP | Synapt G2-S |

After chromatographic separation, the oligonucleotides are ionized using electrospray ionisation in negative mode and enter the mass spectrometer (all five datasets). Subsequent detection was achieved by high-resolution mass spectrometers (HRMS), either an orbitrap (from Thermo Fisher, used for 3 datasets) or a time-of-flight (TOF from Waters, used for 2 datasets) as shown in Figure 1C. All instruments were operated in DDA mode with three datasets acquired using CID in a HCD collision cell or linear ion trap (Thermo Fisher) and two datasets using CID in a Triwave collision cell (Waters). As oligonucleotides fragment at specific bonds (27), the resulting MS/MS spectra are used for sequence assignment.

Data analysis for four workflows used the NucleicAcidSearchEngine (NASE) (15) while System 5 used Pytheas (23) for MS/MS sequence assignments as described in Materials & Methods. The known modifications in 28S rRNA (1) were added to the sequence file for static search mode while a search of the unmodified rRNA sequence using both variable location and variable chemical identity was conducted in dynamic search mode.

### Independent workflows converge on highly similar sequence coverage

With CID, RNA oligonucleotide ions undergo backbone cleavage primarily at one of the phosphodiester bonds and the 2’-C-O bond, generating characteristic fragment ions originating from the 5′ end (a and c ions) and the 3′ end (w and y ions), with the subscript indicating the cleavage position (Figure 2A). Other fragment types (b/x, d/z) can arise via secondary dissociation at higher energies but are typically low in abundance (28). An exemplary MS/MS spectrum of the modified oligonucleotide UACCC[Am]UAUCCGp together with the annotated fragment ions is shown in Figure 2B. These MS/MS spectra are matched to candidate oligonucleotide sequences that can then be used for mapping onto the 28S rRNA sequence. Although instrument architectures and experimental parameters - especially the available collision energy - differed, the systems produced essentially identical fragment ladders, with variations in the relative intensities of the dominant ions (Figure S5).

**Figure 2.**
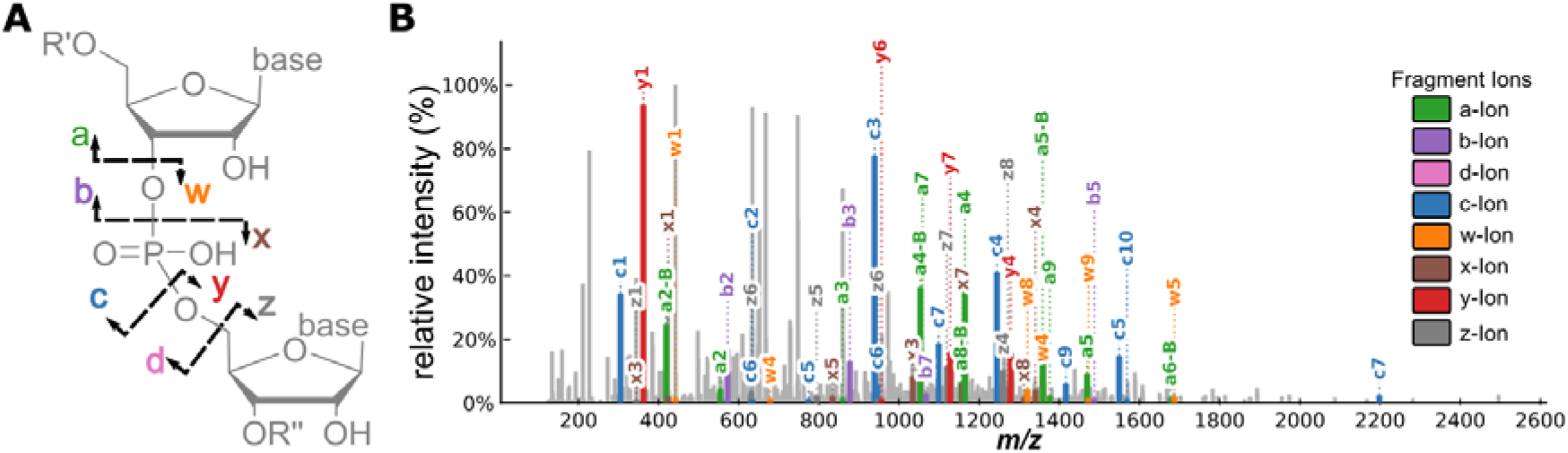
Overview of MS/MS spectra for sequence annotation. **A** Fragmentation of oligonucleotides alongside the phosphodiester and the fragment nomenclature established by McLuckey (27). **B** Exemplary MS/MS spectrum of the [Am]-modified oligonucleotide from the analytical flow-HILIC workflow with Exploris 240 orbitrap detection and HCD fragmentation (System 1). Resulting fragments are colour coded according to the McLuckey definition given in A.

To compare sequence assignment across workflows, all identified oligonucleotides were projected onto the mature human 28S rRNA sequence using a genome browser-style visualization (Figure 3A). In this representation, the x-axis corresponds to the nucleotide position within the mature 28S rRNA, while the y-axis distinguishes uniquely mapped from non-uniquely mapped oligonucleotides. Unique oligonucleotides map to a single position within the reference sequence, whereas non-unique oligonucleotides represent short sequence stretches that occur multiple times within the rRNA and therefore cannot be assigned unambiguously.

**Figure 3.**
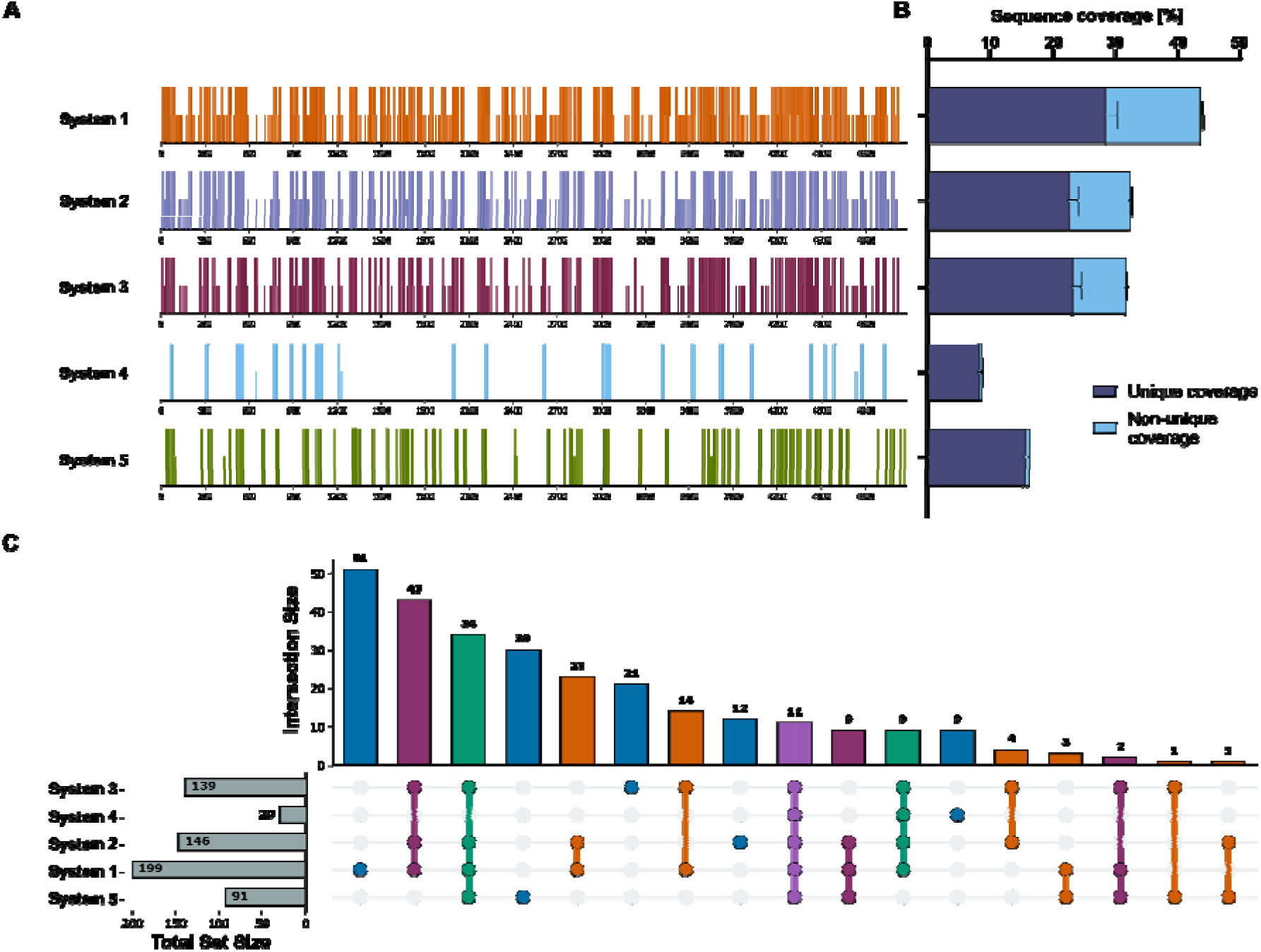
Fragments identified from the human 28S rRNA after a partial RNase T1 digest. **A** Genome browser style display of the human 28S rRNA (MODOMICS ID:160 M11167.1) with the sequence position given on the x-axis and the uniqueness on the y-axis. Half-length bars indicate non-unique sequences, full-length bars indicate unique sequences. **B** Sequence coverage for each workflow as stacked bar graphs (Average of 6 different reference IDs). Dark blue indicates unique sequences, light blue indicates non-unique sequences. **C** Upset plot for comparison of found fragment overlap detected by the different workflows (MODOMICS ID:160 M11167.1).

Sequence coverage was subsequently calculated for each workflow using either only uniquely mapped oligonucleotides or all identified oligonucleotides, including non-unique sequence assignments (Figure 3B). Using uniquely mapped fragments, sequence coverage ranged from approximately 10–28% across the five workflows. Inclusion of non-unique oligonucleotides increased the apparent sequence coverage up to ∼40%. This latter observation is largely a consequence of the digestion strategy rather than analytical performance. Human 28S rRNA is highly GC-rich (∼ 70%) and, following partial RNase T1 digestion, generates numerous short oligonucleotides terminating at guanosine residues. Many of these fragments are too short to be assigned uniquely within the approximately 5 kb long rRNA sequence. Consequently, sequence coverage calculated from all identified oligonucleotides overestimates the amount of uniquely resolved sequence information.

Partial rather than complete RNase T1 digestion generated numerous missed-cleavage products spanning one or more additional guanosine residues. The results demonstrate that controlled partial digestion of a base-specific endonuclease (like RNase T1) is one strategy for maximizing informative sequence coverage in oligonucleotide mass spectrometry. However, longer oligonucleotides also increase the complexity of database searching by substantially expanding the number of possible digestion products. Consequently, partial digestion requires careful optimization to balance gains in unique sequence information against increased computational complexity and search-space expansion.

To quantify the reproducibility of fragment identification across workflows, the overlap of all identified oligonucleotides was assessed using an UpSet plot (Figure 3C). In total, 277 unique oligonucleotides were identified across all participating workflows. Of these, 46 mapped oligonucleotides were identified by two workflows, 54 were identified by three workflows, 43 were identified by four workflows, and 11 were detected in all five workflows. Altogether, 144 unique oligonucleotides (55.6%) were detected by at least two workflows. The remaining 123 unique oligonucleotides (44.4%) were observed by a single workflow.

Importantly, the UpSet analysis demonstrates that workflows with higher sequence coverage did not identify fundamentally different regions of the 28S rRNA but instead recovered additional oligonucleotides from the same underlying sequence space. Conversely, workflows with lower coverage largely represented subsets of the higher-coverage datasets rather than complementary fragment populations. Thus, differences between workflows primarily reflected analytical depth rather than qualitative differences in sequence recovery.

Taken together, these results demonstrate a remarkably high degree of reproducibility across independent LC-MS/MS workflows. Despite differences in chromatographic separation, mass spectrometer architecture, and instrumental implementation, a significant level of sequence information was consistently recovered. This observation indicates that, under controlled sample and harmonized analysis conditions, a variety of LC-MS/MS workflows are robust and transferable analytical frameworks for sequence-resolved RNA analysis.

### Search-parameter choices and precursor isotope handling influence database-search performance

While the LC-MS/MS workflows yielded reproducible data, the automated interpretation of MS/MS data in MS-Seq is another area where additional benchmarking and investigation is warranted. Thus, we focused on using the four datasets that used NASE to better understand what data processing factors impact interlaboratory reproducibility (the dataset from system 5 was excluded as that dataset was analysed using Pytheas). We first tested whether common search-space parameters, including missed-cleavage and sodium-adduct allowances, substantially affected calculated sequence coverage. Allowing between 0 and 10 missed cleavages changed the calculated sequence coverage by less than 3.5 percentage points across all datasets (using static search). Likewise, permitting up to four sodium adducts per precursor altered coverage by less than 4 percentage points. These results indicate that neither missed cleavage allowance nor precursor adducts substantially affects sequence coverage under the evaluated conditions of static search.

We next tested a precursor-specific parameter: the allowed isotope-offset window used to match observed precursor *m/z* values to theoretical oligonucleotide isotope envelopes. When DDA is used for MS/MS, the instrument generally selects the most abundant ion for fragmentation dependent on instrument settings and exclusion rules. Depending on the selection window width, the number of isotopic peaks selected for MS/MS can vary. Further, oligonucleotides with >9 nt will have a base peak (i.e., most abundant isotopic peak) that is not the monoisotopic peak due to the presence of ^13^C,^18^O or ^15^N. Thus, it is important to understand how MS platform software assignments of precursor ions impact downstream data processing.

We tested two precursor isotope-offset windows during database searching. M4 allowed the observed precursor *m/z* to match the theoretical monoisotopic peak through the +4 isotope peak. M5 expanded this window by also allowing a match to the −1 isotope peak, in addition to the monoisotopic through +4 isotope peak. We found that sequence coverage varied minimally between M5 and M4 for system 1 (39.4%→38.9%) and system 3 (29.6%→30.0%) datasets. System 5 sequence coverage decreased slightly between M5 and M4 (10.9%→7.6%) while system 2 decreased significantly between M5 and M4 (31.3%→16.8%). The strong coverage drop for system 2 suggests that many assigned spectra depended on allowing the observed precursor *m/z* to match non-monoisotopic isotope positions within candidate oligonucleotide envelopes. These results strongly suggest that precursor isotope-offset windows can impact downstream data processing and will require the analyst to ensure an appropriate offset is used.

We also investigated whether platform-specific processing (vendor-specific algorithms) also affects the confidence of database identifications. To do so, we compared the NASE Hyperscore for assigned oligonucleotides across the four workflows. The NASE Hyperscore is a spectral-match score reflecting the agreement between an experimental MS/MS spectrum and the corresponding in silico fragment spectrum. Hyperscores for all assigned oligonucleotides (total spectra) were determined. We then removed redundant sequence assignments to determine all non-redundant spectra. Surprisingly, NASE Hyperscore distributions differed markedly between workflows despite comparable numbers of identified oligonucleotides. The dataset from system 2 yielded a median NASE Hyperscore of only 9.4, whereas median scores ranged from 92.1 to 143.0 for the remaining datasets. Nevertheless, system 2 identified 157 non-redundant oligonucleotides, which while below system 1 (222 non-redundant oligonucleotides), is comparable to system 3 (155 non-redundant oligonucleotides) (Supplementary Table S2)

To investigate whether these Hyperscore differences simply reflected differences in the analysed oligonucleotide population, we compared the length distributions of the non-redundant identified sequences. While system 4 contained somewhat longer oligonucleotides on average (18.1 nt), systems 1-3 showed comparable mean sequence lengths between 10.6 and 14.7 nt. Thus, differences in oligonucleotide length cannot explain the significantly lower NASE Hyperscores observed for system 2.

The origin of these Hyperscore differences remains unclear but may reflect differences introduced during vendor-specific precursor detection or peak picking prior to database searching. This observation warrants further investigation but does not affect the overall benchmark conclusions. Importantly, these observations demonstrate that NASE Hyperscores are not necessarily directly comparable across instrument platforms and underline the importance of transparent reporting and harmonization of processing workflows in RNA-MS benchmarking studies.

### Modification-aware database searches dramatically expand the search space

The benchmark analyses presented above were performed using a MODOMICS (1) annotated reference sequence in which both the identity and genomic position of known 28S rRNA modifications were predefined. NASE or Pytheas was then used in static search mode to confirm these previously annotated RNA modifications. However, another important role for RNA sequencing by LC-MS/MS is the discovery of previously unknown modification sites. Such analyses require dynamic searches in which the reference sequence contains only canonical nucleotides with both variable location and variable chemical identity for the RNA modifications.

To mimic a *de novo* modification discovery experiment, all modified nucleotides in the human 28S rRNA reference sequence were replaced by their corresponding canonical nucleotides prior to database searching. Then, using the dataset obtained from system 1, we analysed the data using NASE in dynamic search mode by providing a defined set of known rRNA modifications as variable modifications that could occur at any compatible nucleotide position within an oligonucleotide.

The identification of RNA modifications by mass spectrometry faces significant challenges due to the existence of isomeric species that share identical molecular masses and elemental compositions. Prominent examples include base methylations and 2′-O-methylations (collectively referred to here as mXn). This poses a particular challenge for MS-based oligonucleotide sequencing because current search engines rely predominantly on precursor and fragment ion masses (16).

Another modification that conventional MS-based sequencing cannot distinguish is pseudouridine (Ψ), which is isomeric to uridine. Including Ψ as a variable modification in a dynamic search presents a particular challenge because it is the only isomer of a canonical nucleoside. Consequently, NASE may computationally assign virtually any uridine residue as pseudouridine, artificially increasing the number of candidate identifications.

A further recurring challenge was the presence of incomplete fragment ion ladders, which resulted in ambiguous modification-site assignments. Such incomplete ladders are frequently observed when fragmentation conditions are suboptimal for oligonucleotides of a particular length.

To avoid these challenges, Ψ was excluded from the reference sequence to prevent artificial inflation of the search space and to maintain robust false discovery rate (FDR) control. In contrast, ribose-methylated species were retained as variable modifications because they occur considerably less frequently than canonical uridines, thereby limiting the increase in computational complexity while preserving biological relevance. Consequently, the search space increased with the number of variable modifications permitted per oligonucleotide.

Allowing a single variable modification per oligonucleotide resulted in only a moderate expansion of the search space (Figure 5A). Of the 253 sequences reported by the dynamic search, 160 were also identified by the static search, whereas 93 additional candidate sequences were exclusively generated by the dynamic search. At this level of search-space expansion, most confidently identified oligonucleotides remained shared between both search strategies, indicating that dynamic searches with a single variable modification remain computationally manageable.

**Figure 4.**
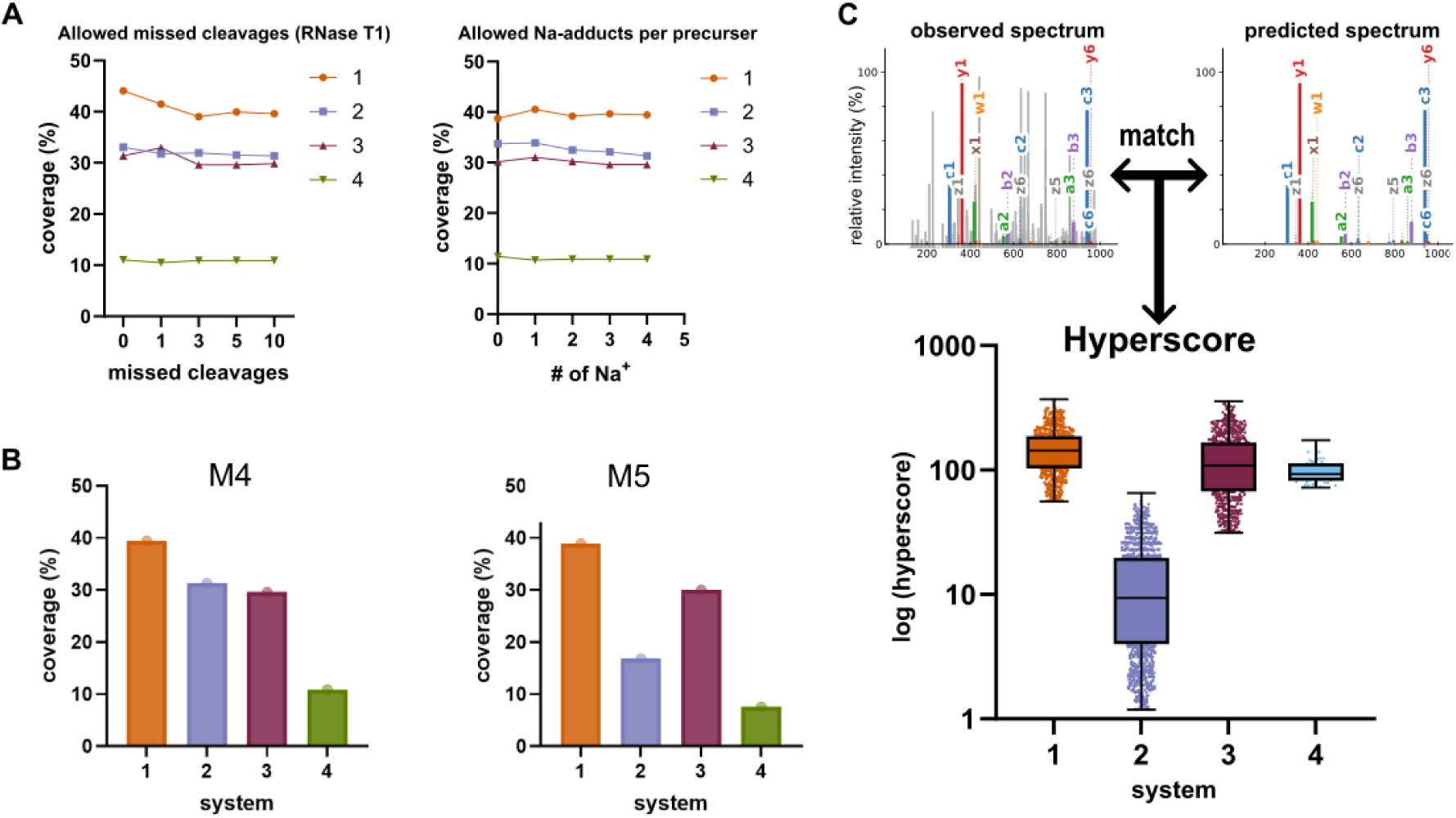
Influence of precursor selection on oligonucleotide identification. **A** Effect of the number of allowed missed RNase T1 cleavages and sodium adducts per precursor on calculated sequence coverage. **B** Effect of precursor isotope envelope-offset on calculated sequence coverage. Restricting precursor selection to the monoisotopic peak markedly reduced coverage for system 2 but had little effect on the remaining platforms. **C** Comparison of NASE Hyperscore distributions and oligonucleotide length distributions across platforms. Despite comparable numbers of non-redundant identified oligonucleotides, system 2 yielded substantially lower NASE scores. Differences in oligonucleotide length did not explain the observed score distributions.

**Figure 5.**
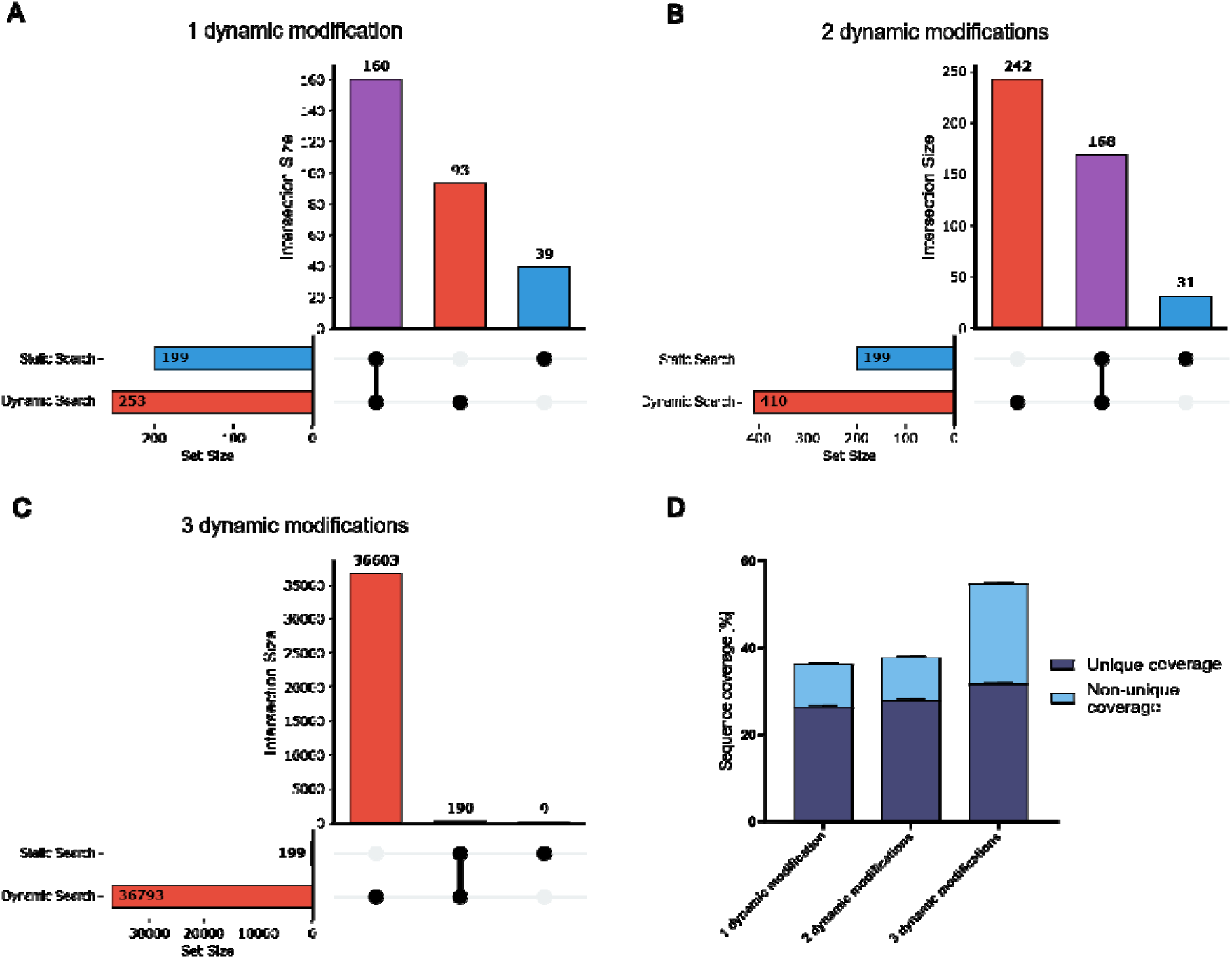
Increasing the number of allowed variable modifications dramatically expands the dynamic search space while only marginally improving sequence coverage. Dynamic searches allowing one (**A**), two (**B**), or three (**C**) variable modifications per oligonucleotide were compared with a static search using a MODOMICS annotated 28S rRNA reference sequence. UpSet plots illustrate the overlap between oligonucleotides identified by both search strategies and the increasing number of dynamic-search-specific candidate assignments. (**D**) Total and unique sequence coverage together with the proportion of ambiguous sequence assignments for the different search strategies.

Increasing the maximum number of variable modifications per oligonucleotide dramatically amplified this effect. Allowing two variable modifications per fragment increased the number of dynamic-search-specific candidate sequences from 93 to 242 (Figure 5B), whereas the number of sequences shared between the static and dynamic searches increased only marginally. Expanding the search further to three variable modifications per oligonucleotide resulted in a combinatorial explosion of the search space. Under these conditions, NASE reported 36,793 candidate sequences that were exclusively identified by the dynamic search, while the number of sequences shared with the static search remained essentially unchanged (Figure 5C).

The consequences of search-space expansion become apparent when comparing the resulting sequence coverage and assignment ambiguity (Figure 5D). Increasing the number of permitted variable modifications only marginally improved both total and unique sequence coverage. In contrast, the proportion of ambiguous sequence assignments increased continuously with search-space expansion. Thus, the large number of additional candidate sequences generated by dynamic searches translated into only minimal gains in informative sequence recovery while substantially increasing the complexity of data interpretation.

These findings demonstrate that comprehensive RNA modification discovery cannot be achieved by unrestricted expansion of the search space alone. Instead, successful *de novo* modification mapping requires carefully optimized search parameters that maximize recovery of correctly identified modification-containing oligonucleotides while minimizing ambiguous candidate assignments. The benchmark presented here was therefore performed using a MODOMICS annotated reference sequence to establish robust search parameters under controlled conditions before extending the analysis to more challenging discovery-oriented search strategies.

## Discussion

### Lessons from benchmarking current oligonucleotide MS workflows

The primary objective of this benchmarking study was not to identify a superior analytical workflow but to evaluate the current capabilities of state-of-the-art oligonucleotide mass spectrometry under standardized conditions. By providing all participating laboratories with an identical ready-to-inject RNA digest and harmonizing data analysis wherever possible, the benchmark deliberately isolated variability arising from chromatographic separation, mass spectrometric detection and instrumental implementation. Under these controlled conditions, five different workflows recovered remarkably similar sets of oligonucleotides (Figure 3) despite substantial differences in chromatographic principles, flow regimes and mass spectrometer architectures. This finding demonstrates that current oligonucleotide LC-MS workflows already provide a robust analytical foundation for sequence-resolved RNA analysis. Importantly, this conclusion should not be interpreted as evidence that current workflows have reached maturity. Rather, the present study establishes an upper estimate of the reproducibility that can currently be achieved under controlled analytical conditions.

### Current limitations in sequence coverage

Despite the encouraging reproducibility observed across workflows, the benchmark also highlights one limitation in this approach. Even under highly standardized analytical conditions, unique sequence coverage remained below one third of the mature 28S rRNA sequence. This observation arises specifically due to the bottom-up approach used here. RNase T1 cleaves specifically after guanosine residues, generating numerous short oligonucleotides from the GC-rich human 28S rRNA. Many of these fragments are too short to be uniquely assigned within the approximately 5 kb transcript and therefore contribute little to sequence coverage despite being confidently identified. Therefore, partial rather than complete RNase T1 digestion was selected as a deliberate strategy to increase sequence information (17,29,30). Missed cleavage events generate longer oligonucleotides that span additional guanosine residues and thereby dramatically increase the probability of unique sequence assignment. This benefit, however, comes at the expense of increased search-space complexity (31), requiring careful optimization of digestion conditions. While there have been several advances in the development of RNases with complementary sequence specificities (12,32–35), additional development of enzymes that can yield longer oligonucleotides and a better understanding of cleavage efficiencies at modified nucleotides remain critical needs for this field.

### Dissecting analytical and biological sources of variability

By distributing an identical ready-to-inject partial RNase T1 digest to all participating laboratories, this benchmark largely isolated analytical variability from biological and pre-analytical sources, highlighting the value of standardized reference materials for objective method comparison. Such reference materials are well established in other omics disciplines (18), where they have enabled objective comparison of analytical workflows, facilitated method development and supported inter-laboratory harmonization. Comparable resources remain largely unavailable for RNA oligonucleotide mass spectrometry. Establishing community reference materials together with harmonized data-analysis strategies should therefore become a priority for future benchmarking efforts and for the continued development of reproducible MS-Seq workflows. However, even under these highly controlled conditions, accurate identification of modified oligonucleotides remained dependent on carefully optimized database-search strategies. The benchmark therefore also highlights that future improvements in MS-Seq will require advances not only in experimental workflows but equally in computational approaches for confident RNA modification discovery.

### Challenges of de novo RNA modification discovery

While the present benchmark demonstrates that current LC-MS workflows can reproducibly recover large parts of the detectable oligonucleotide population, comprehensive identification of RNA modifications remains considerably more challenging. Static database searches using reference sequences with known modifications annotated in place are well suited to confirm known modification patterns. However, MS-Seq can contribute even greater biological insights through the discovery of previously unknown or condition-dependent RNA modifications.

Our comparison of static and dynamic database searches demonstrates that de novo modification discovery is fundamentally limited by combinatorial search-space expansion. Increasing the number of permitted variable modifications generated orders of magnitude more candidate assignments while yielding only marginal improvements in informative sequence coverage. The principal challenge therefore no longer lies in generating additional candidate sequences, but in confidently distinguishing biologically correct assignments from the rapidly growing number of mathematically plausible alternatives. Looking ahead, approaches established in the proteomics field may help improve false discovery rate estimation for RNA oligonucleotide identification in a robust and scalable manner (36).

### Community Consensus and Roadmap for Advancing MS-based sequencing

The observations made during this benchmark closely mirrored the topics discussed during the HRPC satellite meeting. Although participating laboratories employed diverse analytical platforms and data-analysis strategies, there was broad consensus that the next phase of MS-Seq development will depend less on incremental improvements of individual workflows than on coordinated community efforts. Participants identified common priorities spanning instrumentation, RNA chemistry, bioinformatics, metadata standards and reference resources, forming the basis of the community roadmap presented below. Three main areas were discussed: instrumentation, RNA chemistry, and bioinformatics and community resources. The corresponding bottlenecks and recommended workflow steps are summarized in Tables 2 and 3. Based on these findings and the outcomes of the HRPC satellite meeting, we recommend the field focus on the following priorities (Supplemental Table S3) to advance MS-Seq:

1. Establish community reference materials
2. Develop benchmark datasets and reporting standards
3. Build shared MS-Seq resources
4. Advance *de novo* modification bioinformatics

**Table 2:**
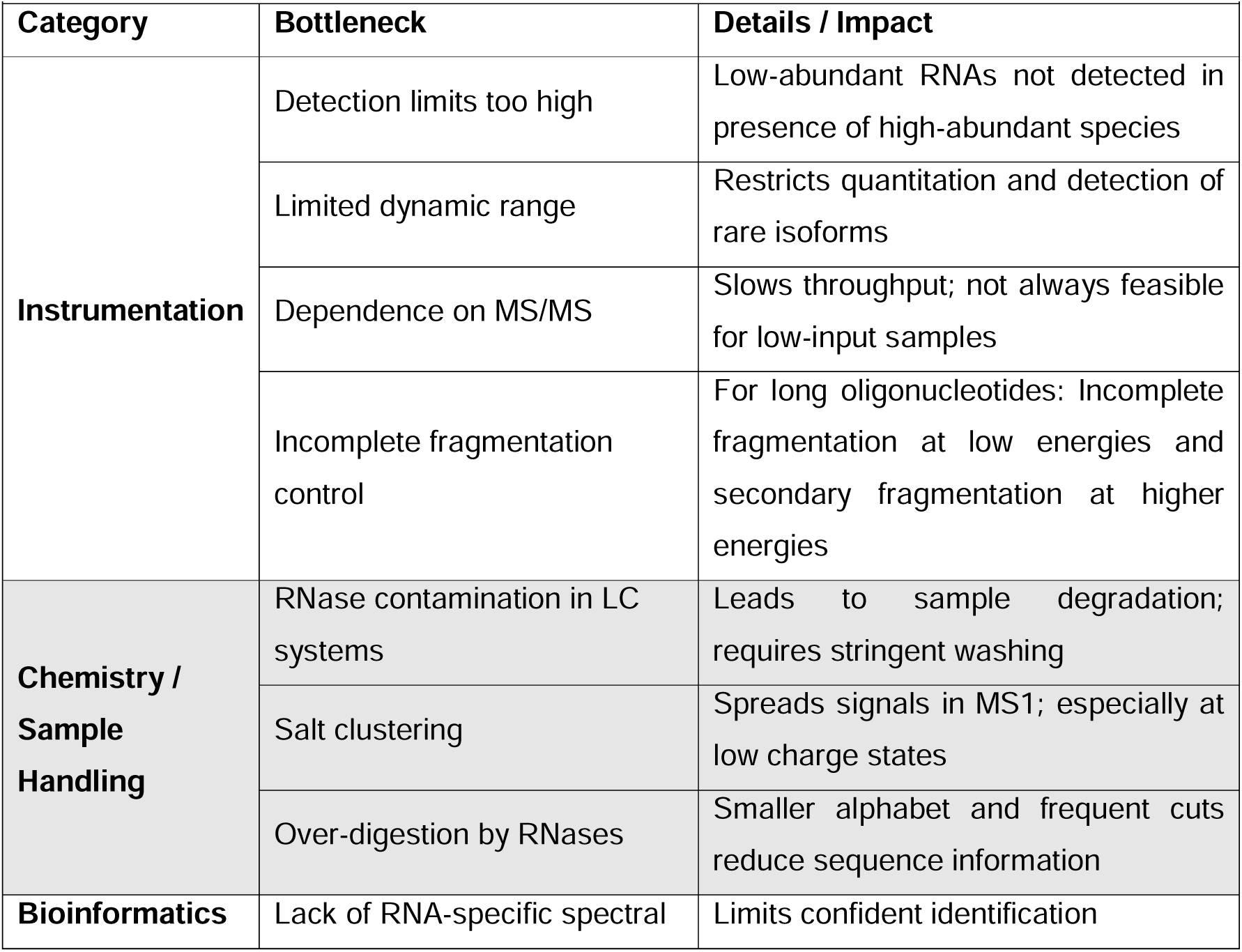
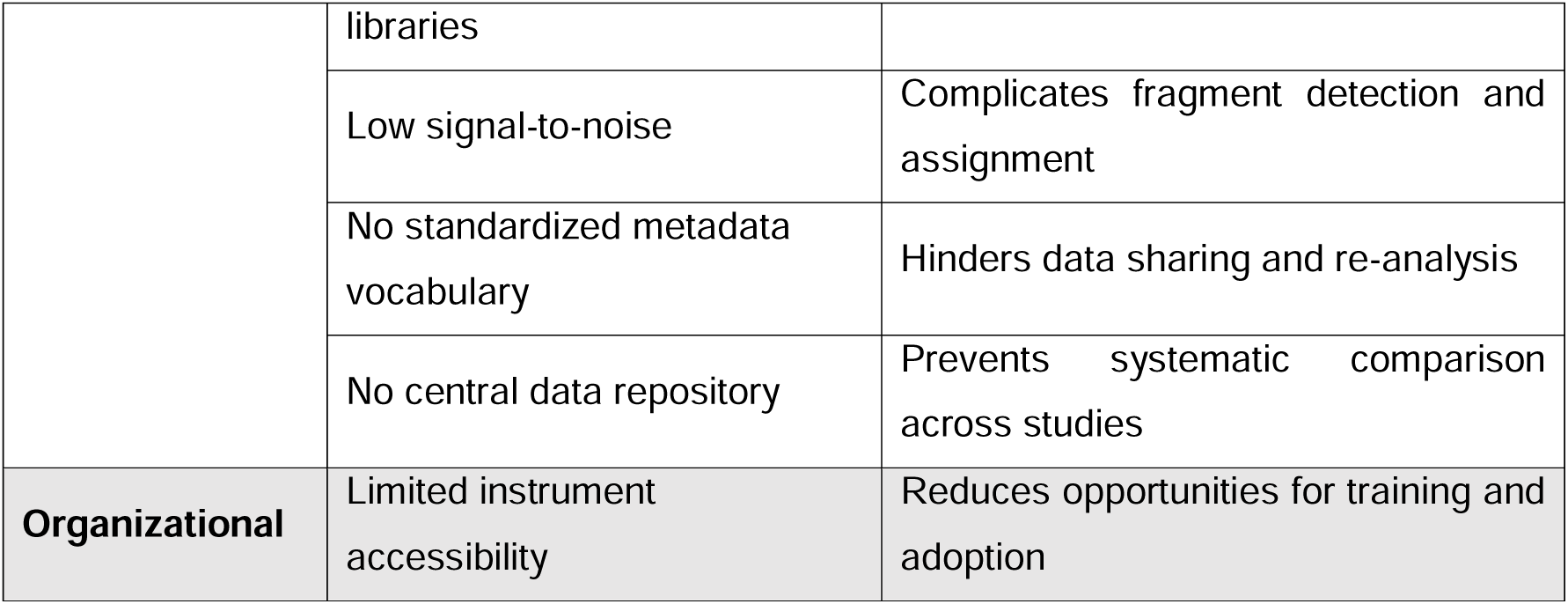
Bottlenecks of current oligonucleotide-MS approaches.

**Table 3:** Critical steps within the LC-MS workflow.

| Workflow Stage | Critical step | Rationale / outcome |
| --- | --- | --- |
| **Sample Preparation** | Fractionation prior to analysis | Increases signal intensity and notability |
|  | Proper desalting | Prevents salt clusters, improves sensitivity |
|  | Controlled RNase digestion | Balances cleavage frequency with information content |
| **Chromatography** | Selecting appropriate chemistry (ion-pairing vs. HILIC) | Impacts separation quality, ionization efficiency |
|  | Preventing RNase carry-over | Ensures reproducibility and avoids degradation |
| **Mass Spectrometry** | MS-only sequencing capability | Reduces sample requirement and acquisition time |
| **Data Analysis** | Implementing FDR with decoys | Ensures confidence in identifications |
|  | Establishing spectral libraries | Improves search performance and comparability |
|  | Standardizing metadata vocabulary | Facilitates data integration |
| **Collaboration** | Sharing raw data in a central repository | Enables re-analysis and benchmarking |
|  | Coordinated reporting of coverage, IDs, modifications | Supports cross-platform comparison |

## Conclusion

This benchmark study demonstrates that current oligonucleotide LC-MS/MS workflows already generate reproducible sequence information across laboratories when common reference materials and harmonized analysis strategies are used. However, substantial challenges remain in coverage, modification discovery and data interoperability. Addressing these challenges will require coordinated community efforts in instrumentation, chemistry, bioinformatics and data standardization.

## Supporting information

Metadata Sheet

## Acknowledgements

This work originated from the 2025 Human RNome Project Consortium (HRPC) Workshop held in Frankfurt, Germany. The workshop was made possible through the generous support of the Warren Alpert Foundation, Brown University, Goethe University Frankfurt, Freunde der Goethe-Universität, the Deutsche Pharmazeutische Gesellschaft (DPhG), House of Pharma, the Collaborative Research Centre SFB 1309, the RNA Modification and Processing (RMaP) initiative, Bruker, Thermo Fisher Scientific, Alida Biosciences, and Biospring. Their support enabled the international gathering of the consortium and the collaborative benchmarking study presented here. The authors further thank all workshop participants for their scientific contributions, open discussions, and collaborative spirit, which laid the foundation for this consortium study. We gratefully acknowledge time and resources invested by our industry partners Thermo Fisher Scientific and Bruker Daltonics GmbH & Co. KG in support of the workshop. We thank Birgit Bartussek for her administrative and organizational contribution to the workshop and all former and current members of the Kaiser/Kellner lab.

## Data availability

The data supporting this article have been included as part of the Supplementary Information.

