## Supplemental Information for "Human 28S rRNA analysed by state-of-the-art oligonucleotide mass spectrometry: benchmarking current capabilities and a call to action for MS-Seq"

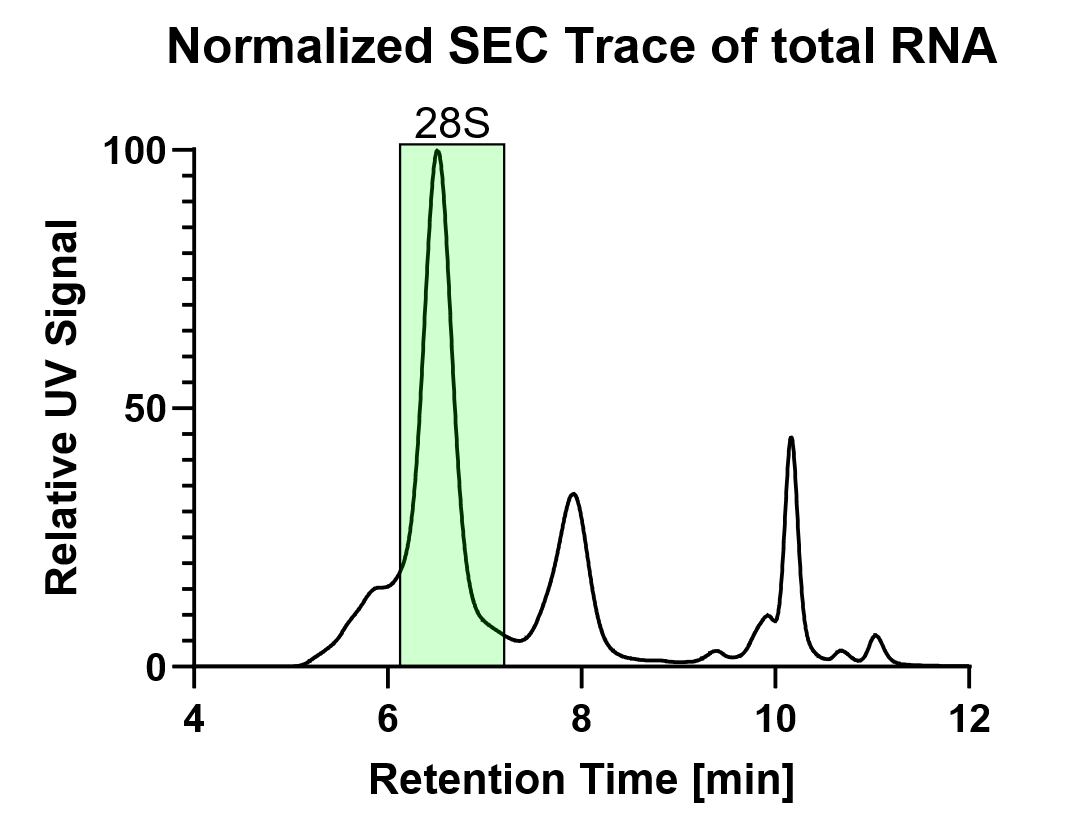

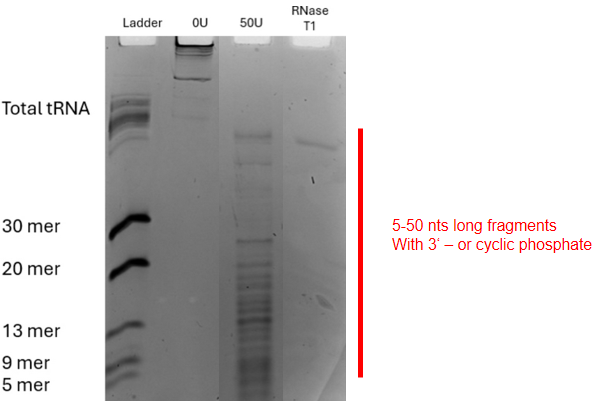

**Figure S1** HEK 293 total RNA was size purified using SEC prior to RNase T1 cleavage of 28S rRNA. **Left:** Normalized SEC trace of total RNA separated on an Agilent HPLC 1200 Series with an Agilent Bio SEC-5 1000Å column. The 28S rRNA fraction collected is shown in the green box and ranges from 6.1 – 7.35 minutes. **Right:** 320 µg 28S rRNA was digested with RNase T1 to yield fragments from 5-50 nts. Enzymes and salts were removed using Zymo Oligo Clean&Concentrator kit. Aliquots of 25 µg were aliquoted and are stored at -80 °C.

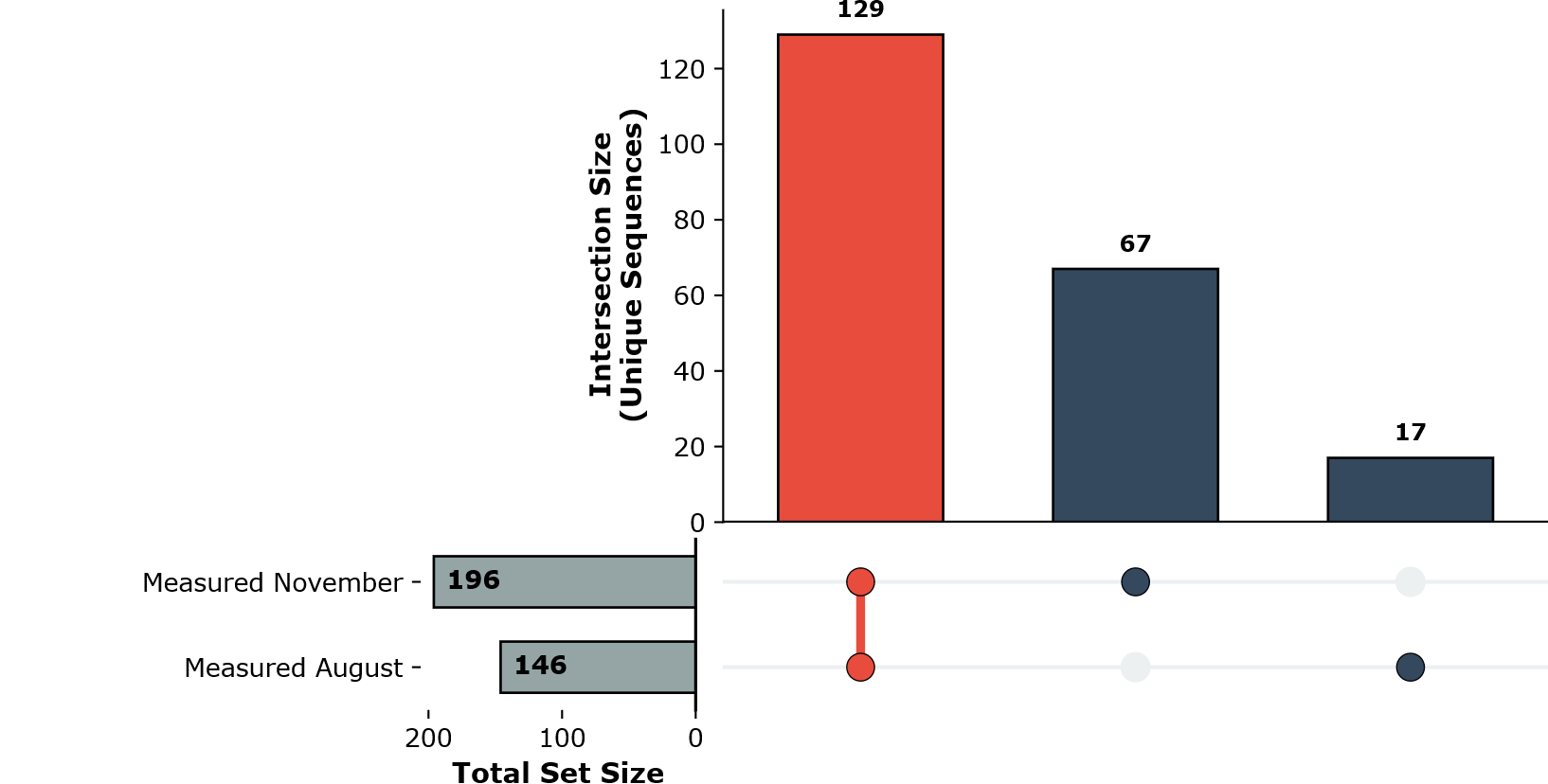

**Figure S2** Comparison of the same 28S rRNA digest measured directly after digestion and after 4 months of storage at -20°C (system 2) (Modomics ID:160 M11167.1).

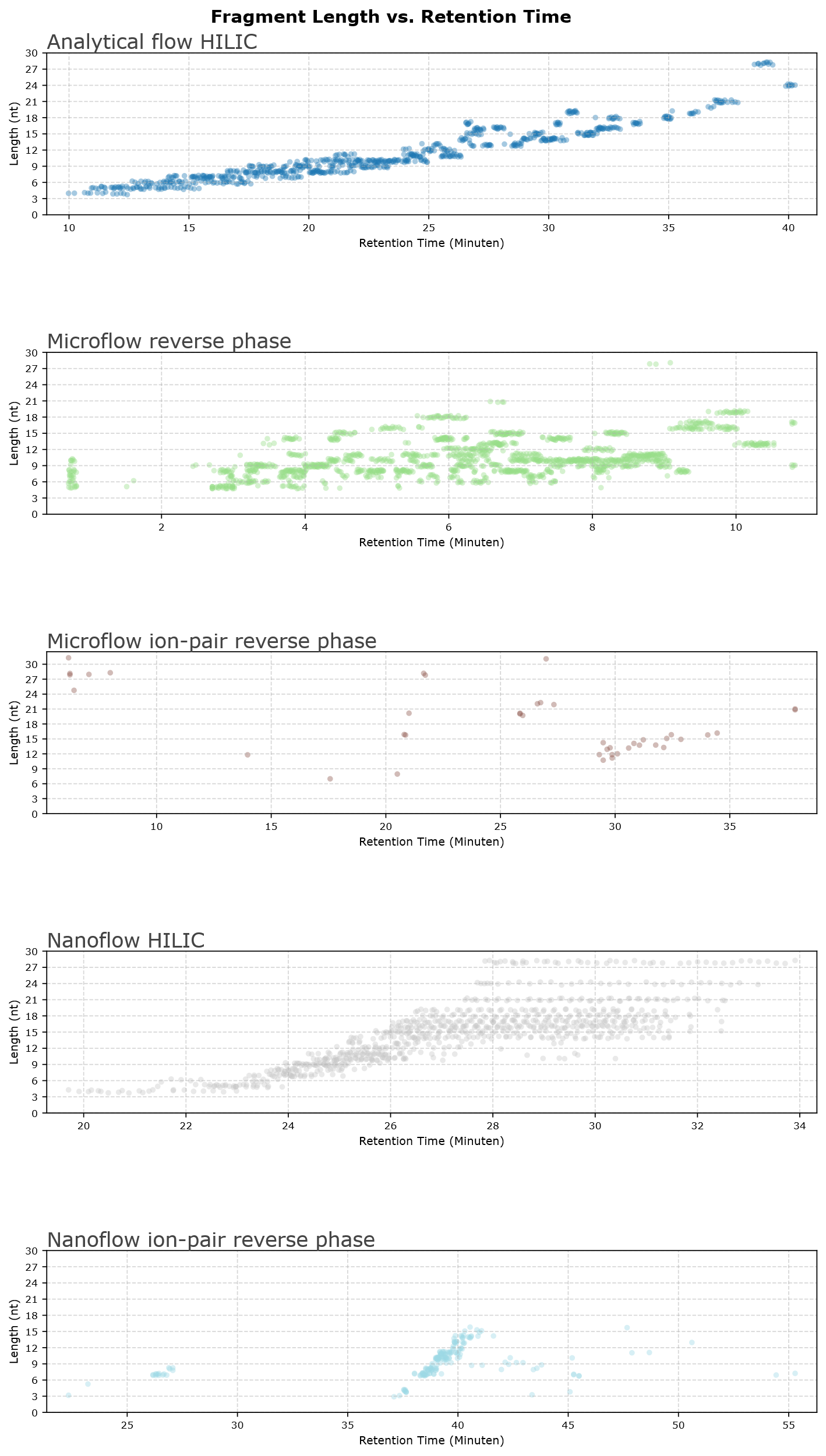

**Figure S3** Length distribution of detected oligonucleotides found throughout the chromatographic window for **A** analytical flow HILIC, **B** microflow reverse phase, **C** micro flow ion-pair reverse phase, **D** nanoflow HILIC, **E** nanoflow ion-pair reverse phase

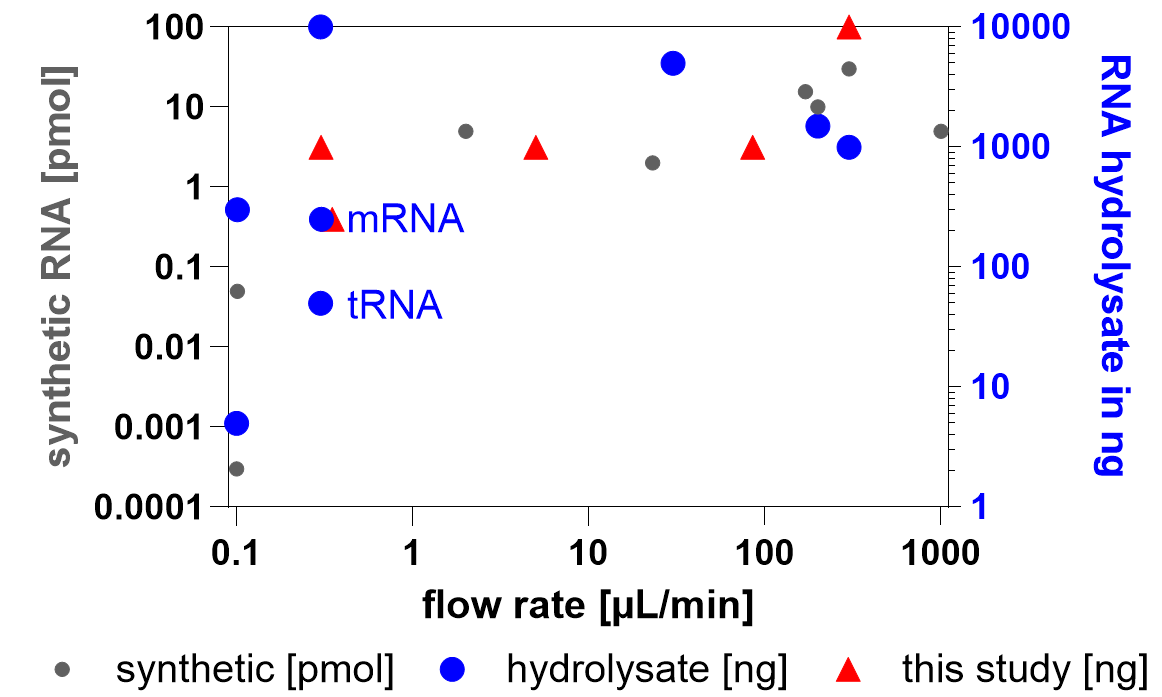

**Figure S4** Sensitivity of reported oligonucleotide MS methods including the injection amounts used in this study (red triangle).

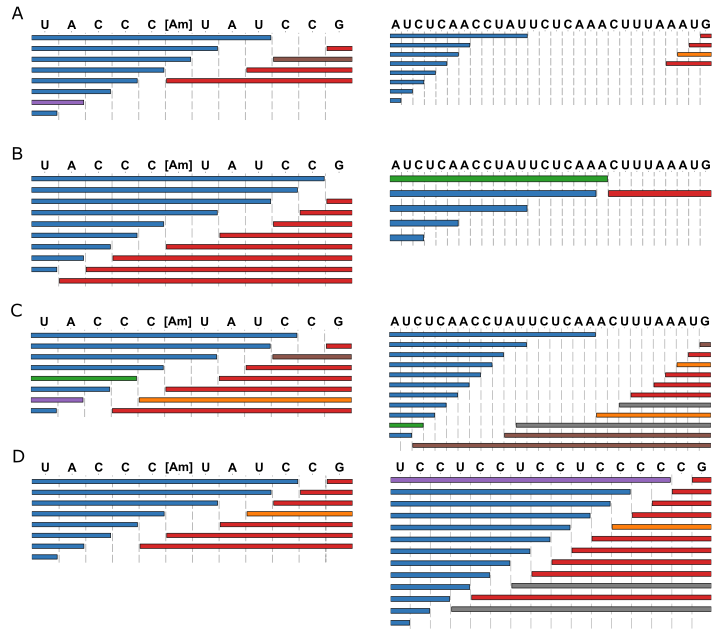

**Figure S5** **Overview of MS/MS spectra and differences among platforms.** **A-D** Fragment map ladder showing only the most intense ion per cleavage site. The sequence is indicated above the respective ladder. Fragments assigned to peaks with less than 5% intensity are not shown. [Am] indicates 2’-O-methyladenosine. **A** System 1 HCD (analytical-HILIC, Exploris 240) **B** System 2 (platform microflow-RP, Eclipse with ion trap CID) **C** System 3 HCD (platform nanoflow-HILIC, q Exactive Plus) **D** System 5 CID (nanoflow IP-RP, Synapt G2-S)

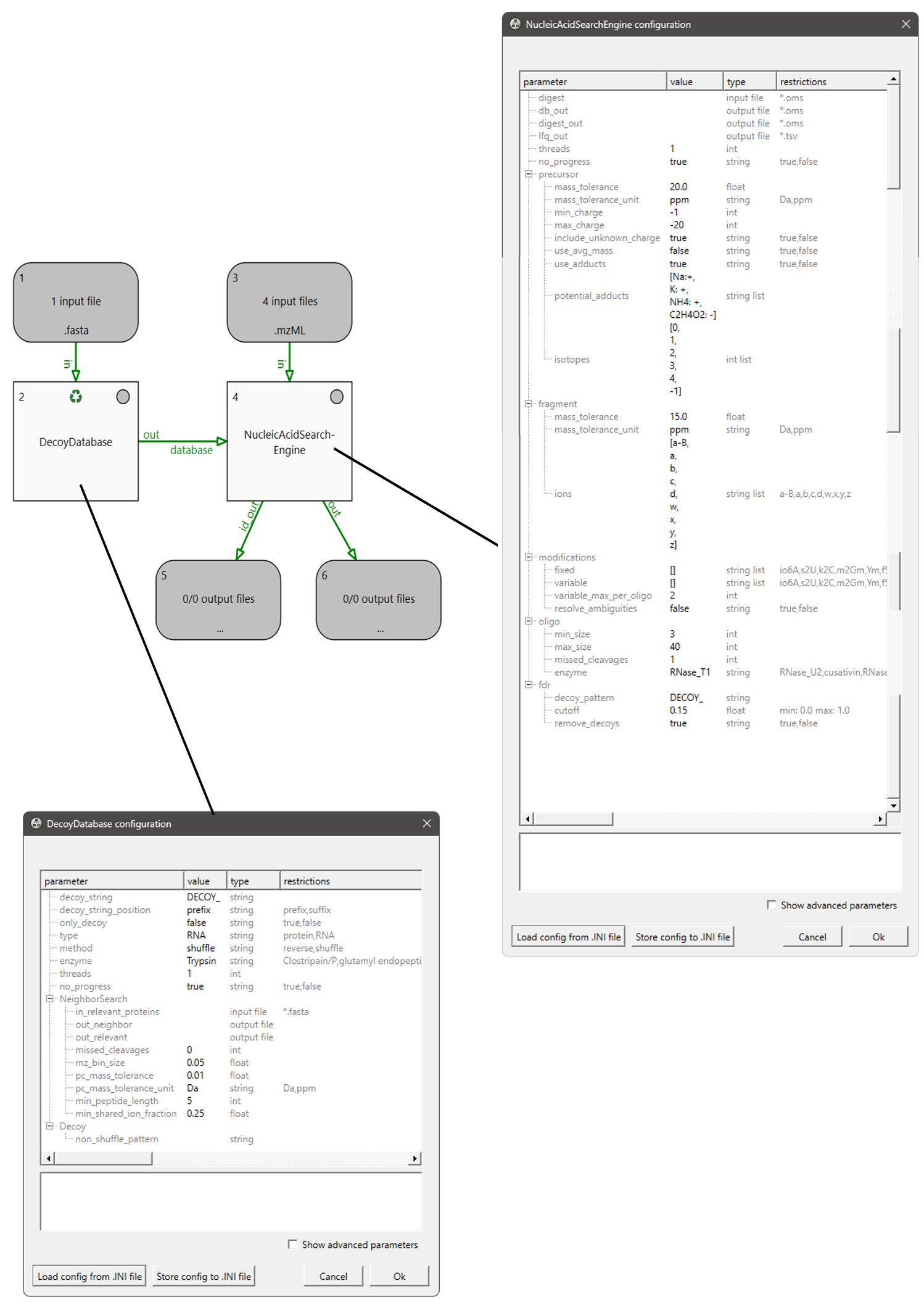

**Figure S6** Overview of NASE parameters and setup for the static search. The settings for the NucleicAcidSearchEngine are seen on the right. While on the bottom left all parameters for decoy generation are shown.

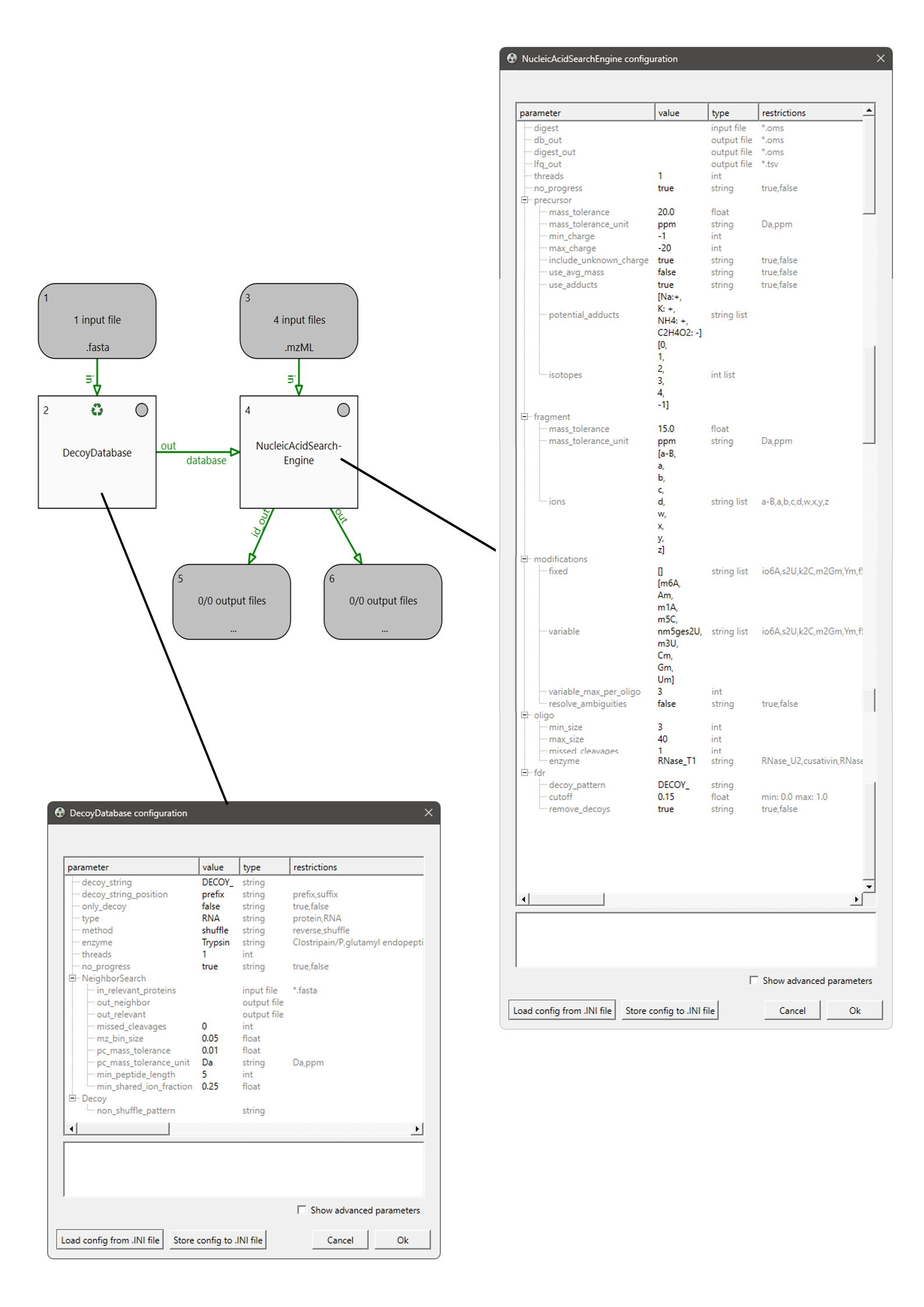

**Figure S7** Overview of NASE parameters and setup for the dynamic search. The settings for the NucleicAcidSearchEngine are seen on the right. While on the bottom left all parameters for decoy generation are shown.

| **Table S1 Metadata collected during the Workshop** | | | | |
| --- | --- | --- | --- | --- |
| **sample name** | **digestion conditions** | **injection** | **Chromatograohy method** | **Mass spec method** |
|  | **date of digestion** | **method reference** | **LC (model, manufacturer, reference, serial number)** | **MS (model, manufacturer, serial number)** |
|  | **amount digested [µg]** | **Sample type (Standard, digest, blank)** | **autosample (model)** | **source type** |
|  | **enzyme (type, vendor, batch, conc, conc-unit)** | **replicate** | **MPA (composition)** | **emitter (type, diameter)** |
|  | **buffer (composition, pH)** | **injected volume** | **MPB (composition)** | **voltage** |
|  | **temperature [°C]** | **amount [pmol/ng]** | **flow rate** | **source temperature** |
|  | **digestion time** |  | **column parameters (type, length, diameter, particle size)** | **reference** |
|  | **additives (type, conc)** |  | **gradient** | **aquisition type (DDA,DIA,SRM etc.)** |
|  | **replicate** |  |  | **aquisition method (reference)** |

**Table S2:** Fragment information extracted from NASE mzTab results. The table shows total spectra identified, non-redundand sequences identified, NASE hyperscore and length information.

| **NASE Version** | **Sample Name** | **Source File** | **Total Spectra (OSM)** | **Unique Sequences (OSM)** |
| --- | --- | --- | --- | --- |
| 3.5.0 | System 1 | 02_28SrRNA_2.mzTab | 790 | 222 |
| 3.5.0 | System 2 | 20250810_rRNA_1ug_A002_031.mzTab | 1450 | 157 |
| 3.5.0 | System 4 | 20260213_rRNA_1ug_70%B_upto_70min.mzTab | 42 | 34 |
| 3.5.0 | System 3 | B02_004_28S_250ng28S_1pmolSynth_MPB_  SCID30_HCD30.mzTab | 854 | 155 |
| - | System 5 | final_report_G02_007_rRNA_1000ng (2).csv | - | - |

| **Sample Name** | **Score Min** | **Score Max** | **Score Mean** | **Score Median** | **Length Min** | **Length Max** | **Length Mean** |
| --- | --- | --- | --- | --- | --- | --- | --- |
| System 1 | 55.64 | 366.35 | 151.09 | 142.99 | 4 | 28 | 11.22 |
| System 2 | 1.19 | 64.59 | 13.49 | 9.37 | 5 | 28 | 10.59 |
| System 4 | 72.35 | 170.64 | 99.07 | 92.05 | 7 | 31 | 18.12 |
| System 3 | 31.33 | 352.57 | 122.98 | 108.78 | 4 | 28 | 14.72 |
| System 5 | - | - | - | - | - | - |  |

**Table S3 Community consensus priorities for advancing MS-based RNA sequencing (MS-Seq)**

| **Area** | **Current consensus** | **Priority actions** |
| --- | --- | --- |
| **Instrumentation** | Current LC-MS platforms already provide reproducible sequence information across laboratories. Remaining limitations primarily concern sensitivity, dynamic range and low-abundance RNAs. | Improve sensitivity and dynamic range; develop electron-based fragmentation, ion mobility and MS-only sequencing strategies; enable routine analysis of longer oligonucleotides. |
| **RNA chemistry** | RNA sample preparation strongly influences informative sequence recovery. Partial digestion, desalting and chromatographic separation remain major determinants of unique sequence coverage. | Develop complementary RNases with alternative cleavage specificities; characterize RNase cleavage efficiencies at modified nucleotides; improve chromatographic workflows and sample preparation. |
| **Bioinformatics** | Database searching becomes increasingly ambiguous with expanding modification search space. Harmonized workflows are required for reproducible analyses. | Develop modification-aware search algorithms, improved FDR estimation, ambiguity-aware scoring strategies and de novo modification discovery approaches. |
| **Reference materials** | Benchmarking requires common biological reference materials to separate analytical from biological variability. | Establish standardized RNA reference materials and defined digestion standards for community-wide benchmarking. |
| **Metadata and reporting** | Reproducibility depends on complete reporting of experimental and computational parameters. | Define minimum metadata requirements, standardized coverage definitions, reporting guidelines and controlled vocabularies. |
| **Community resources** | Public infrastructure for RNA-MS data remains limited. | Establish community repositories, spectral libraries and FAIR-compliant reference datasets. |
| **Human RNome Project** | Different biological questions require different analytical priorities. | Prioritize sequence coverage for sequence confirmation, chromatographic resolution for impurity characterization, and both high coverage and chromatographic performance for discovery-driven studies. |
